# UV and bacteriophages as a chemical-free approach for cleaning membranes from anaerobic bioreactors

**DOI:** 10.1101/2020.08.03.234450

**Authors:** Giantommaso Scarascia, Luca Fortunato, Yevhen Myshkevych, Hong Cheng, TorOve Leiknes, Pei-Ying Hong

## Abstract

Anaerobic membrane bioreactor (AnMBR) for wastewater treatment has attracted much interest due to its efficacy in providing high quality effluent with minimal energy costs. However, membrane biofouling represents the main bottleneck for AnMBR because it diminishes flux and necessitates frequent replacement of membranes. In this study, we assessed the feasibility of combining bacteriophages and UV-C irradiation to provide a chemical-free approach to remove biofoulants on the membrane. The combination of bacteriophage and UV-C resulted in better log cells removal and twice higher extracellular polymeric substance (EPS) concentration reduction in mature biofoulants compared to UV-C. A reduction in the relative abundance of *Acinetobacter* spp. and selected gram-positive bacteria associated with the membrane biofilm was also achieved by the new cleaning approach. Microscopic analysis further revealed the formation of cavities in the biofilm due to bacteriophages and UV-C irradiation, which would be beneficial to maintain water flux through the membrane. When the combined treatment was further compared with the common chemical cleaning procedure, a similar reduction on the cell numbers was observed (1.4 log). However, combined treatment was less effective in removing EPS compared with chemical cleaning. These results suggest that the combination of UV-C and bacteriophage have an additive effect in biofouling reduction, representing a potential chemical-free method to remove reversible biofoulants on membrane fitted in an anaerobic membrane bioreactor.

**SIGNIFICANCE:** Anaerobic membrane bioreactors can achieve high quality effluent with a reduced energy consumption. However, biofouling represents the main bottleneck for membrane filtration efficiency. Biofouling is commonly reduced through chemical treatment. These agents are often detrimental for the environment and health safety due to the formation of toxic byproducts. Therefore, we present a new approach, based on the additive antifouling action of bacteriophages infection and UV-C irradiation, to reduce anaerobic membrane biofouling. This new strategy could potentially delay the occurrence of membrane fouling by removing the reversible fouling layers on membranes, in turn reducing the frequencies and amount of chemicals needed throughout the course of wastewater treatment.

## INTRODUCTION

Anaerobic membrane bioreactor (AnMBR) has emerged as an energy-efficient biotechnology for municipal wastewater treatment (1). The post-AnMBR effluent has low turbidity, retains the ammonium and phosphate, and hence can potentially be reused as liquid fertilizers for agricultural crops (2). Moreover, AnMBR offers numerous advantages. Anaerobic mode of wastewater treatment eliminates the need for aeration, hence saving up to 75% of the energy costs associated with conventional wastewater treatment (3). The organic carbon in wastewater is fermented to produce methane, which can be combusted and converted to electrical energy. Furthermore, anaerobic process results in lower sludge production than aerobic treatment, therefore minimizing solid waste disposal costs (1, 4). Finally, the coupling of membrane to anaerobic fermentation tanks reduces the required footprint for wastewater treatment process by eliminating the need for clarifiers (5, 6).

Despite these advantages, AnMBR is prone to membrane biofouling due to the unwanted deposition of microorganisms and their extracellular polymeric substances (EPS) on the membrane surface (7). This deposition of biofoulants decreases the operational flux and rapidly increases the transmembrane pressure (TMP) and energy required to maintain a constant flux (8). Strategies to control biofouling in AnMBR can be devised based on our understanding of the main biofoulant constituents. The AnMBR biofilm microbial community was recently revealed to assemble based on random stochastic events. However, a few core genera (e.g. *Methanobacterium* and *Acinetobacter*) occurred in relative abundance that deviated from the neutral assembly model (9). This suggests that they may be playing a potential keystone role in the formation of biofoulant layers on the membrane and can be potential targets to eliminate in a bid to alleviate membrane biofouling.

Current strategies to reduce membrane biofouling includes the use of physical means (e.g. gas scouring, backwashing and sonication) and chemical cleaning (e.g. citric acid and chlorine (5)). Physical cleaning is generally more effective for the reversible foulant layer (10), while chemical cleaning is used to remove irreversible foulant layers (11). However, the use of these chemicals can lead to the formation of carcinogenic and toxic byproducts, and can detrimentally impact the membrane integrity (12, 13). In addition, it was shown that certain bacteria, such as *Acinetobacter* spp. are strongly resistant to chemical disinfectants (14).

Biological-based approaches have emerged as interesting alternatives to chemical-based methods of alleviating membrane biofouling (15, 16). In particular, bacteriophages are viruses that infect viable bacteria and have been used against biofilm-associated bacteria for clinical therapeutic treatments (17–19). Besides infecting their intended bacterial host, bacteriophages can also induce the release of enzymes that degrade the EPS biofilm matrix, in turn, increasing biofilm susceptibility to biocides (20). Moreover, bacteriophages are able to infect bacteria over a broad range of pH, salinity and temperature (21). For these reasons, bacteriophages were also applied as a biological-based approach to reduce membrane biofouling (22), albeit mainly demonstrated against single-species biofilms (23, 24). However, the use of bacteriophages alone does not lead to adequate efficacy in terms of biofouling reduction (21) and their effectiveness against a complex biofilm has not been demonstrated. Instead, bacteriophages were applied in combination with chemicals to enhance their effect on membrane biofilm (21, 25). This counteracts the use of biological-based approaches to minimize chemical usage.

To reduce the need of chemicals for membrane cleaning, we applied bacteriophages in combination with ultraviolet (UV-C) irradiation. UV-C (254 nm) imposes germicidal effect by causing DNA damage and it has been widely used for effluent disinfection (26) and for wastewater pretreatment (27). Although UV effect on biofilm was investigated earlier, these studies were dedicated to determine its antibiofilm action within the water distribution system (28, 29) and not for the purpose of mitigating membrane biofouling. By applying bacteriophages together with UV-C irradiation to tackle membrane biofouling, we hypothesize that these two agents can have an additive action against the biofilm matrix. This is because UV-C irradiation was determined previously to trigger bacteriophages to enter into lytic mode and lyse planktonic bacterial cells (30), but such demonstrations on the use of UV-C to enhance the efficiency of bacteriophage against the biofilm matrix have not been conducted.

In this study, we aim to demonstrate the use of bacteriophages in combination with UV-C to reduce membrane biofouling in AnMBR. A cocktail of three isolated *Acinetobacter* spp. bacteriophages was applied in combination with UV-C exposure, and its effect on the membrane biofouling layer was analyzed and compared against individual treatment with bacteriophages or UV-C alone. The treatment combining bacteriophages and UV-C was also compared against the chemical cleaning method to examine its feasibility as an alternative biological-based approach to reduce AnMBR biofouling.

## RESULTS

### Isolated bacteriophages and their ability to infect membrane biofilm

Three isolated *Acinetobacter* bacteriophages were characterized based on their morphology. All three bacteriophages showed regular icosahedral head with no visible tail. The size of the head was ~86 nm for *A. junii phage* (Figure S1A), ~75 nm for *A. modestus* phage (Figure S1B) and ~68 nm for *A. seohaensis* phage (Figure S1C). Based on the Ackermann classification (31), the three bacteriophages isolated in this study could tentatively be placed in the order *Caudovirales*, under the *Podoviridae* family, constituted by an icosahedral head and short or absent tail.

The cocktail of *A. modestus, A junii* and *A. seohaensis* bacteriophages were introduced either alone or in combination with UV-C to the fouled membranes. All three bacteriophages, in the absence of UV-C, increased in their PFUs compared to the initial spiked amount when introduced to membranes harvested at 40 and 60 kPa (Figure S2, p < 0.04). However, only *A. modestus* bacteriophages were able to actively propagate when introduced to membranes harvested at 20 kPa (Figure S2B, p = 0.03), while the rest of the phages did not have PFUs that were statistically different from the initial spiked value (Figure S2A and S2C). In contrast, UV-C + bacteriophages treatment did not result in any increase in PFU numbers of both *A. junii* and *A. seohaensis* bacteriophages for all membranes harvested at 20, 40 and 60 kPa (Figure S2A and S2C). Only *A. modestus* phages increased in its PFU despite the presence of UV-C, and the final PFU numbers were significantly higher compared with the initial value spiked to the membranes harvested at 40 and 60 kPa (Figure S2B, p < 0.001). There was no significant difference in PFU numbers for *A. modestus* phages when spiked to membranes harvested at 20 kPa (p = 0.55). No bacteriophages were recovered after UV-C treatment alone and in the control condition.

### Biofilm analysis: bacterial cells

Compared with the control, the bacteriophages cocktail alone significantly reduced the number of cells associated with the biofilm from 40 kPa membrane (Figure 1A, p = 0.02). The same reduction did not occur for membranes harvested at 20 and 60 kPa (p > 0.15). Both UV-C and UV-C + bacteriophages treatment significantly reduced the number of cells compared to the control for membranes harvested at all biofouling rates analyzed in this study (Figure 1A, p < 0.0001). In particular, UV-C + bacteriophage treatment was able to outcompete the UV-C only treatment for membranes harvested at 40 and 60 kPa (Figure 1A, p < 0.0001), with the UV-C + bacteriophage treatment reporting up to 1.36-log reduction of cells compared with the control. However, there was no statistical difference between the UV-C + bacteriophage treatment and the UV-C only treatment for membranes harvested at 20 kPa (Figure 1A, p = 0.99).

**Figure 1.**
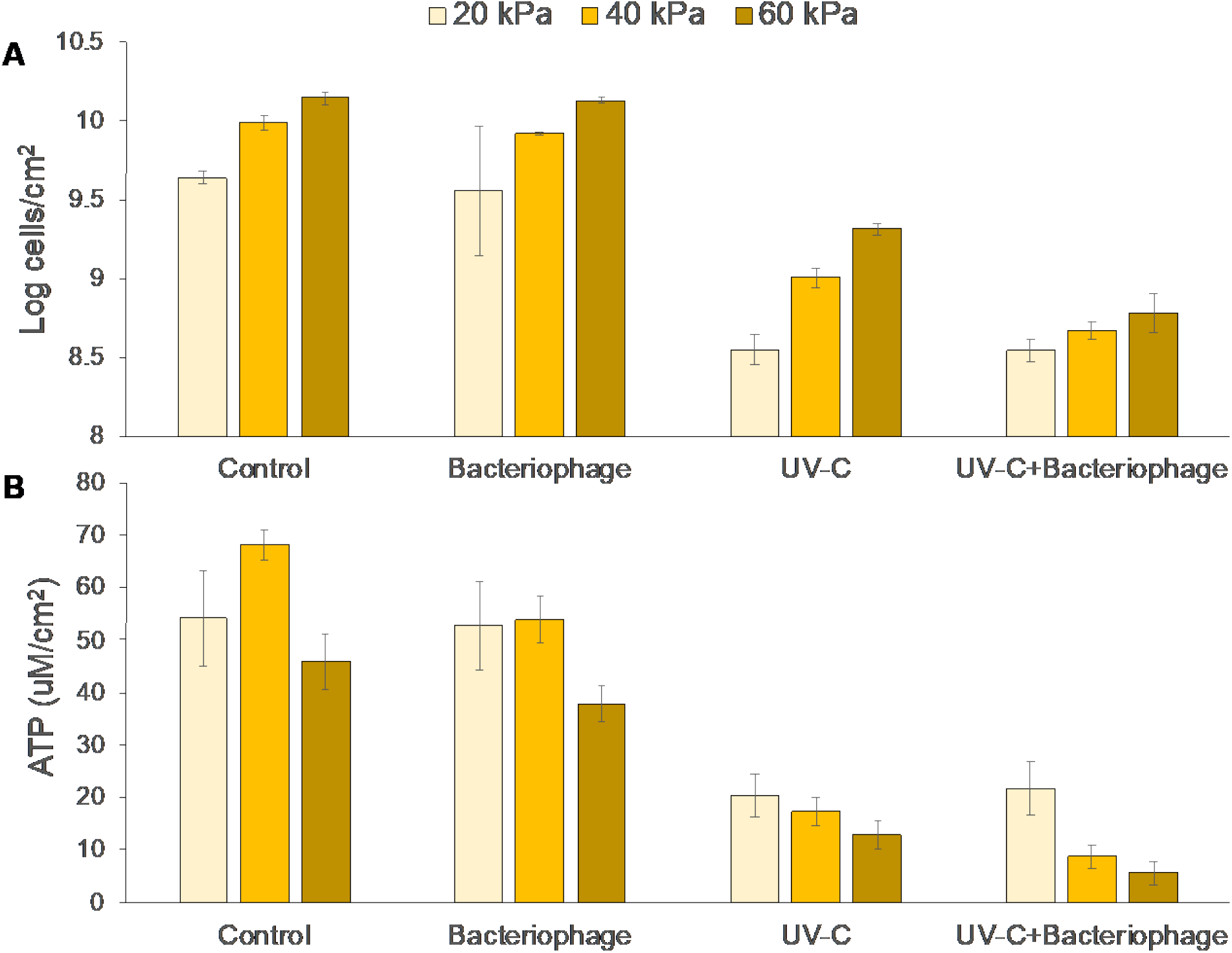
Log of the number of cells (A) and ATP concentration (B) in the membrane biofilm after each treatment. Bar colors indicate different membrane biofouling degree. Error bars indicate standard deviation among the three replicates

Similar trends were observed for the ratio between alive and dead cells in the biofilm. For membranes harvested at 40 and 60 kPa, bacteriophages cocktail (p < 0.01), UV-C only (p < 0.0001) and UV-C + bacteriophage (p < 0.0001) were able to reduce the alive to dead cell ratio compared to the control (Table 1). UV-C + bacteriophage had a significantly lower ratio compared to UV-C only for membranes harvested at 40 and 60 kPa (p < 0.0001), while there was no difference between these two treatments for membranes harvested at 20 kPa (Table 1, p = 0.66). Moreover, there was no difference between control and bacteriophages treatment (p = 0.47) for membrane harvested at 20 kPa while the UV-C + bacteriophage and UV-C only treatments had the same significant effect on the alive to dead cells ratio compared to the control (Table 1, p < 0.0001).

**Table 1.** Ratio between alive and dead cells in membrane biofilm after different treatments and at different biofouling rates. The data are presented as average (n = 3) ± standard deviation.

| Condition | 20 kPa | 40 kPa | 60 kPa |
| --- | --- | --- | --- |
| Control | $1.22 \pm 0.11$ | $1.05 \pm 0.02$ | $1.15 \pm 0.10$ |
| Bacteriophage | $1.17 \pm 0.13$ | $0.84 \pm 0.05$ | $1.01 \pm 0.10$ |
| UV-C | $0.5 \pm 0.09$ | $0.48 \pm 0.08$ | $0.56 \pm 0.08$ |
| UV-C + Bacteriophage | $0.53 \pm 0.14$ | $0.24 \pm 0.05$ | $0.25 \pm 0.09$ |

### Biofilm analysis: ATP

ATP concentration within the biofilm matrix was used as an indicator of cell activity. The maximum cell activity for the control was observed for membranes harvested at 40 kPa (Figure 1B). For membranes harvested at 40 kPa and 60 kPa, ATP concentration was significantly reduced by 14 and 9 μM/cm^2^, respectively, due to bacteriophages application compared to the control (Figure 1B, p < 0.0009). This difference was however not observed for membranes harvested at 20 kPa (p = 0.74). UV-C and UV-C + bacteriophages reduced ATP concentration for all membranes compared with the control (Figure 1B, p < 0.0001). Specifically, UV-C + bacteriophages outcompeted UV-C treatment alone when the membrane was harvested at 40 and 60 kPa (Figure 1B, p < 0.003), achieving a total reduction of at least 50 μM/cm^2^ compared with the control.

### Biofilm analysis: proteins and polysaccharides in biofilm matrix

All treatments (i.e., bacteriophages, UV-C and UV-C + bacteriophage) caused a reduction in both biofilm-associated proteins and polysaccharides concentration on membranes harvested at 40 and 60 kPa compared to control (Figure 2, p < 0.0002). The bacteriophage treatment did not result in any significant reduction in both proteins and polysaccharides concentration on membranes harvested at 20 kPa (p = 0.37). In contrast, UV-C and UV-C + bacteriophages treatments reduced at least 50% of proteins and 35% of polysaccharides concentrations compared with the control across all tested membranes (Figure 2). For membranes harvested at 40 and 60 kPa, UV-C + bacteriophages treatment achieved a statistically higher proteins and polysaccharides reduction compared with UV-C only treatment (Figure 2, p < 0.004), while the difference was not significant for the membranes harvested at 20 kPa (p > 0.24).

**Figure 2.**
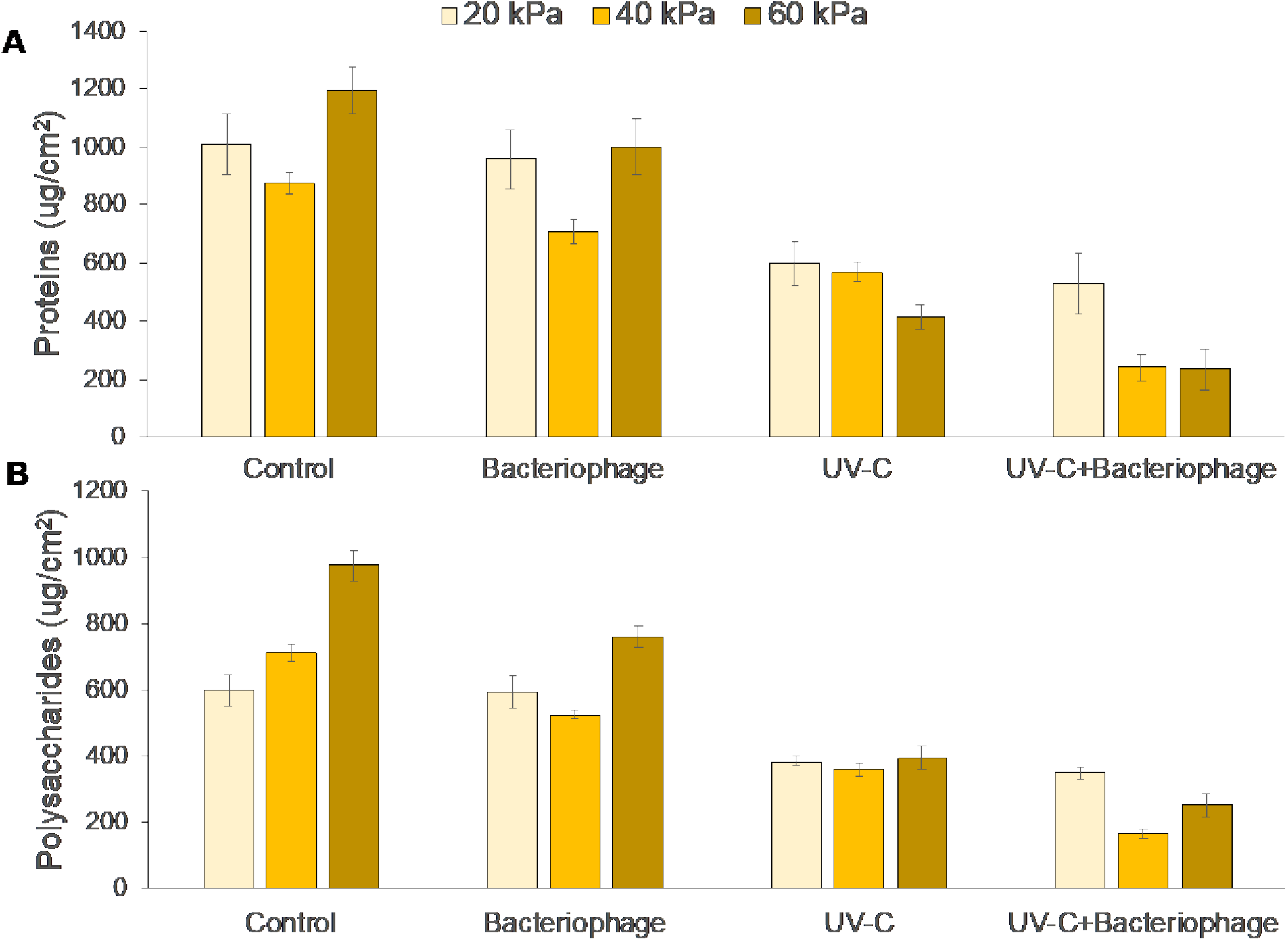
Proteins (A) and polysaccharides (B) concentration in the membrane biofilm after each treatment. Bar colors indicate different membrane biofouling degree. Error bars indicate standard deviation among the three replicates

### Active bacterial community characterization

Genus *Acinetobacter* was the intended target of the isolated bacteriophages and its relative abundance was reduced significantly across all tested membranes compared with the control when bacteriophages were applied alone or in combination with UV-C (Table 2). Although not the intended target of the isolated bacteriophages, other genera were also affected in their relative abundance. For example, the relative abundance of genus *Pseudomonas* was significantly lower upon bacteriophages and UV-C + bacteriophages treatments but increased in its relative abundance upon UV-C only treatment (Table 2). Moreover, two anaerobic gram-negative genera *Paludibacter* and *Cloacibacterium* also reduced in their relative abundance upon bacteriophages and UV-C + bacteriophage treatment. A similar observation was made for gram-positive bacteria which had a lower relative abundance compared to control upon exposure to bacteriophage and UV-C + bacteriophage (Table 2). UV-C treatment, however, resulted in an increase in the relative abundance for most gram-positive bacteria compared to the control, particularly for membranes harvested at 40 kPa.

**Table 2.** Percentage relative abundance for different genera in control and upon exposure to treatment. Only genera that changed in their relative abundance for all tested membranes were shown. Green and red cells indicate statistically lower and higher relative abundance, respectively, compared to the control (p < 0.05). Stars indicate genera that include gram-positive bacteria. Data are presented as average of biological replicates (n = 3) ± standard deviation.

| Genus | Control | Bacteriophages | UV-C | UV-C + Bacteriophages |
| --- | --- | --- | --- | --- |
| **20 kPa** |  |  |  |  |
| *Micrococcus* * | 0.03 ± 0.02 | 0.00 ± 0.00 | 0.04 ± 0.04 | 0.00 ± 0.00 |
| *Cloacibacterium* | 0.09 ± 0.02 | 0.01 ± 0.00 | 0.10 ± 0.02 | 0.01 ± 0.00 |
| *Paludibacter* | 0.40 ± 0.08 | 0.22 ± 0.14 | 0.54 ± 0.11 | 0.40 ± 0.22 |
| *Acinetobacter* | 6.04 ± 0.94 | 0.01 ± 0.01 | 7.58 ± 0.31 | 0.02 ± 0.01 |
| *Pseudomonas* | 4.67 ± 1.32 | 0.44 ± 0.17 | 5.27 ± 1.42 | 0.50 ± 0.21 |
| *unclassified_Firmicutes* * | 1.56 ± 0.71 | 0.41 ± 0.50 | 1.49 ± 0.92 | 0.57 ± 0.45 |
| *unclassified_Clostridiales* * | 0.12 ± 0.02 | 0.07 ± 0.01 | 0.13 ± 0.04 | 0.08 ± 0.03 |
| *unclassified_Clostridiaceae* * | 0.03 ± 0.01 | 0.02 ± 0.01 | 0.04 ± 0.02 | 0.02 ± 0.01 |
| *Clostridium sensu stricto* * | 0.05 ± 0.03 | 0.02 ± 0.01 | 0.09 ± 0.03 | 0.02 ± 0.01 |
| **40 kPa** |  |  |  |  |
| *Micrococcus* * | 0.04 ± 0.00 | 0.00 ± 0.00 | 0.08 ± 0.02 | 0.00 ± 0.00 |
| *Cloacibacterium* | 0.10 ± 0.01 | 0.01 ± 0.00 | 0.11 ± 0.03 | 0.01 ± 0.01 |
| *Paludibacter* | 0.42 ± 0.02 | 0.22 ± 0.14 | 0.51 ± 0.11 | 0.13 ± 0.00 |
| *Acinetobacter* | 6.99 ± 0.12 | 0.01 ± 0.01 | 7.89 ± 0.63 | 0.00 ± 0.00 |
| *Pseudomonas* | 5.07 ± 0.49 | 0.52 ± 0.08 | 6.20 ± 0.45 | 0.52 ± 0.05 |
| *unclassified_Firmicutes* * | 1.61 ± 0.10 | 0.36 ± 0.45 | 1.91 ± 0.09 | 0.16 ± 0.01 |
| *unclassified_Clostridiales* * | 0.13 ± 0.01 | 0.06 ± 0.01 | 0.17 ± 0.02 | 0.06 ± 0.00 |
| *unclassified_Clostridiaceae* * | 0.05 ± 0.01 | 0.02 ± 0.00 | 0.07 ± 0.01 | 0.02 ± 0.00 |
| *Clostridium sensu stricto* * | 0.06 ± 0.01 | 0.02 ± 0.00 | 0.10 ± 0.03 | 0.03 ± 0.00 |
| **60 kPa** |  |  |  |  |
| *Micrococcus* * | 0.04 ± 0.01 | 0.00 ± 0.00 | 0.06 ± 0.00 | 0.00 ± 0.00 |
| *Cloacibacterium* | 0.10 ± 0.01 | 0.00 ± 0.00 | 0.11 ± 0.01 | 0.01 ± 0.00 |
| *Paludibacter* | 0.43 ± 0.04 | 0.13 ± 0.02 | 0.50 ± 0.10 | 0.26 ± 0.28 |
| *Acinetobacter* | 7.13 ± 0.32 | 0.00 ± 0.00 | 7.85 ± 0.18 | 0.01 ± 0.02 |
| *Pseudomonas* | 5.22 ± 0.18 | 0.51 ± 0.05 | 5.93 ± 0.46 | 0.52 ± 0.24 |
| *unclassified_Firmicutes* * | 1.79 ± 0.10 | 0.13 ± 0.03 | 2.03 ± 0.18 | 0.38 ± 0.44 |
| *unclassified_Clostridiales* * | 0.12 ± 0.00 | 0.07 ± 0.01 | 0.15 ± 0.01 | 0.07 ± 0.02 |
| *unclassified_Clostridiaceae* * | 0.04 ± 0.01 | 0.02 ± 0.00 | 0.05 ± 0.01 | 0.02 ± 0.01 |
| *Clostridium sensu stricto* * | 0.06 ± 0.00 | 0.02 ± 0.00 | 0.06 ± 0.01 | 0.03 ± 0.01 |

### OCT analysis

In-situ non-invasive biomass analysis was accomplished by using OCT on the membrane biofilm portions collected after the experiments to assess the efficiency of fouling mitigation strategy (32). Figure 3 shows an upper view of the 3D cross-sections of the different treatments employed in this study. There was an observed change in the biofilm coverage across the membrane. For example, control sample presented a flat and compact biomass structure that was homogeneously distributed on the membrane surface (Figure 3A). At the end of the exposure, all the treatments led to a partial reduction of the biofilm coverage on the membrane and a more irregular morphology in terms of thickness and roughness (Figure 3 and Table S1). There was at least 11.5% reduction in biofilm coverage compared to the control as a result of the treatments. In addition, a decrease of 20% of biovolume was observed after 6 h exposure to all the treatments compared with the control (Table S1, p < 0.0001).

**Figure 3.**
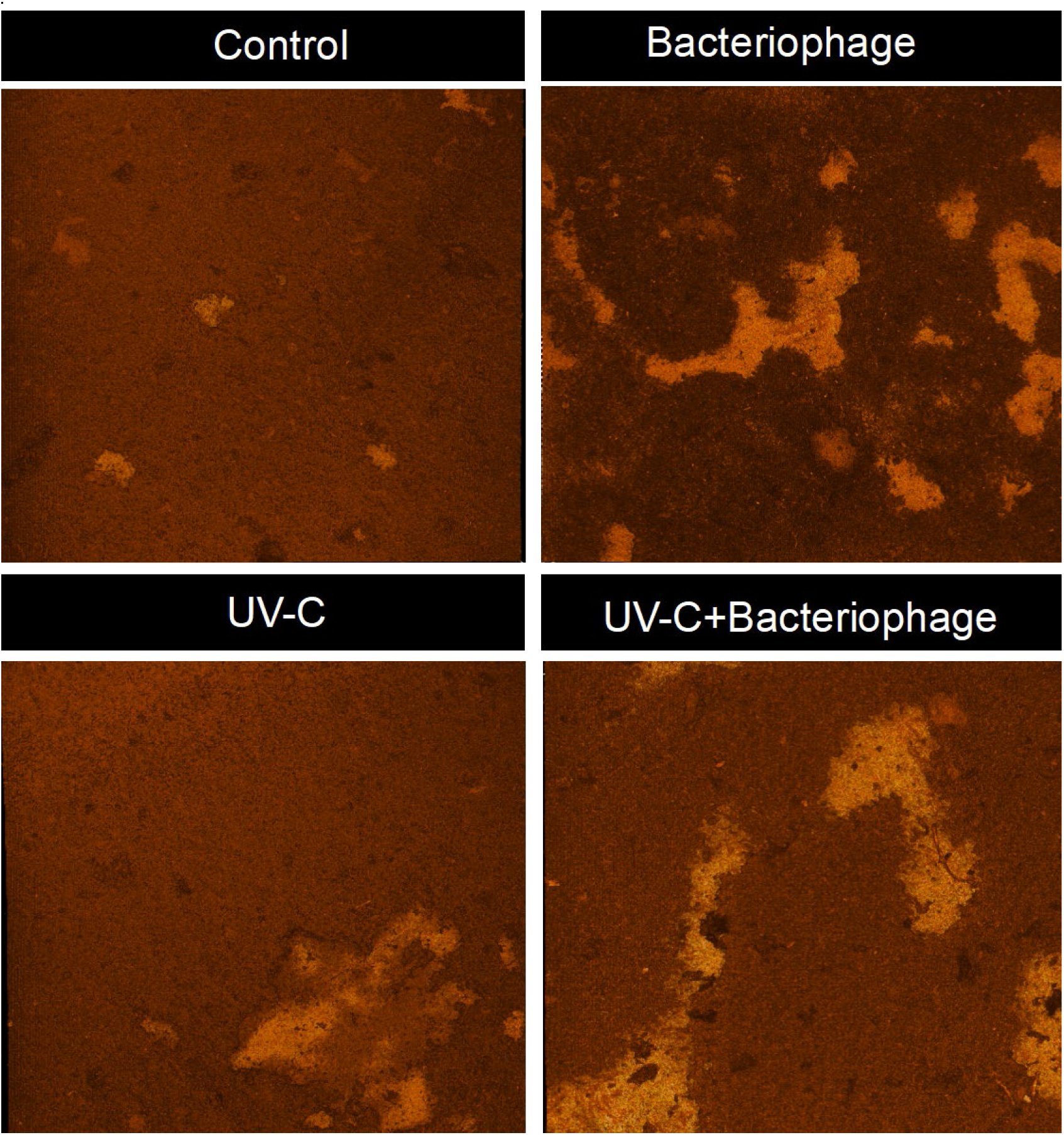
Upper view of 3D OCT scans of the membrane biofilm for control and the three treatment. Darker color indicates thicker biofilm while brighter color represents areas not covered by biofilm. The area visualized is 8 × 8 mm.

Bacteriophage infection alone was further monitored over time in terms of biofilm removal. It was observed that a substantial decrease in biovolume and increase in roughness was observed already after 1 h from the beginning of the bacteriophage treatment (Table S2). A further decrease in biovolume and increase in roughness was observed after 3 h, albeit at a lower extent than that observed in the first hour (Table S2).

### Effect of the order of application of UV-C and bacteriophages

UV-C was applied independently before bacteriophages addition to assess if this order of application will be equally effective in biofilm removal. This order of applying UV-C and bacteriophage is subsequently referred to as “independent UV-C + bacteriophages” treatment. It was observed that UV-C + bacteriophages treatment resulted in statistically higher cells removal (Figure S3A, p = 0.01) and polysaccharides concentration reduction (Figure S3B, p = 0.02) compared to the independent UV-C + bacteriophage treatment. However, there was no statistical difference in the reduction of protein concentration (p = 0.72) achieved by either treatment.

### Comparison with chemical treatment

UV-C + bacteriophage treatment outcompeted the other tested treatments (i.e., bacteriophage and UV-C only), and was therefore further compared against the common chemical (i.e., sodium hypochlorite and citric acid) cleaning process. Both UV-C + bacteriophage and chemical treatment significantly reduced the values of all the biofilm parameters examined, namely cells number, ATP concentration, alive and dead cells ratio, protein and polysaccharides concentration, compared to the control (p < 0.0001, Figure 4 and S4). However, ATP concentration was not significantly different between UV-C + bacteriophage and chemical treatment (p = 0.35). Log cells reduction compared with the control was slightly different between chemical treatment and UV-C + bacteriophage treatment (1.7 and 1.4, respectively, Figure 4A, p = 0.04). Moreover, the value of the ratio of alive and dead cells upon chemical treatment was 0.2, while the one after UV-C + bacteriophage treatment was 0.4 (Figure S4, p = 0.02). In addition, the chemical treatment method was 4 to 6-times more effective in reducing proteins and polysaccharides concentration in biofilm matrix compared to UV-C + bacteriophage treatment (Figure 4B, p < 0.0001).

**Figure 4.**
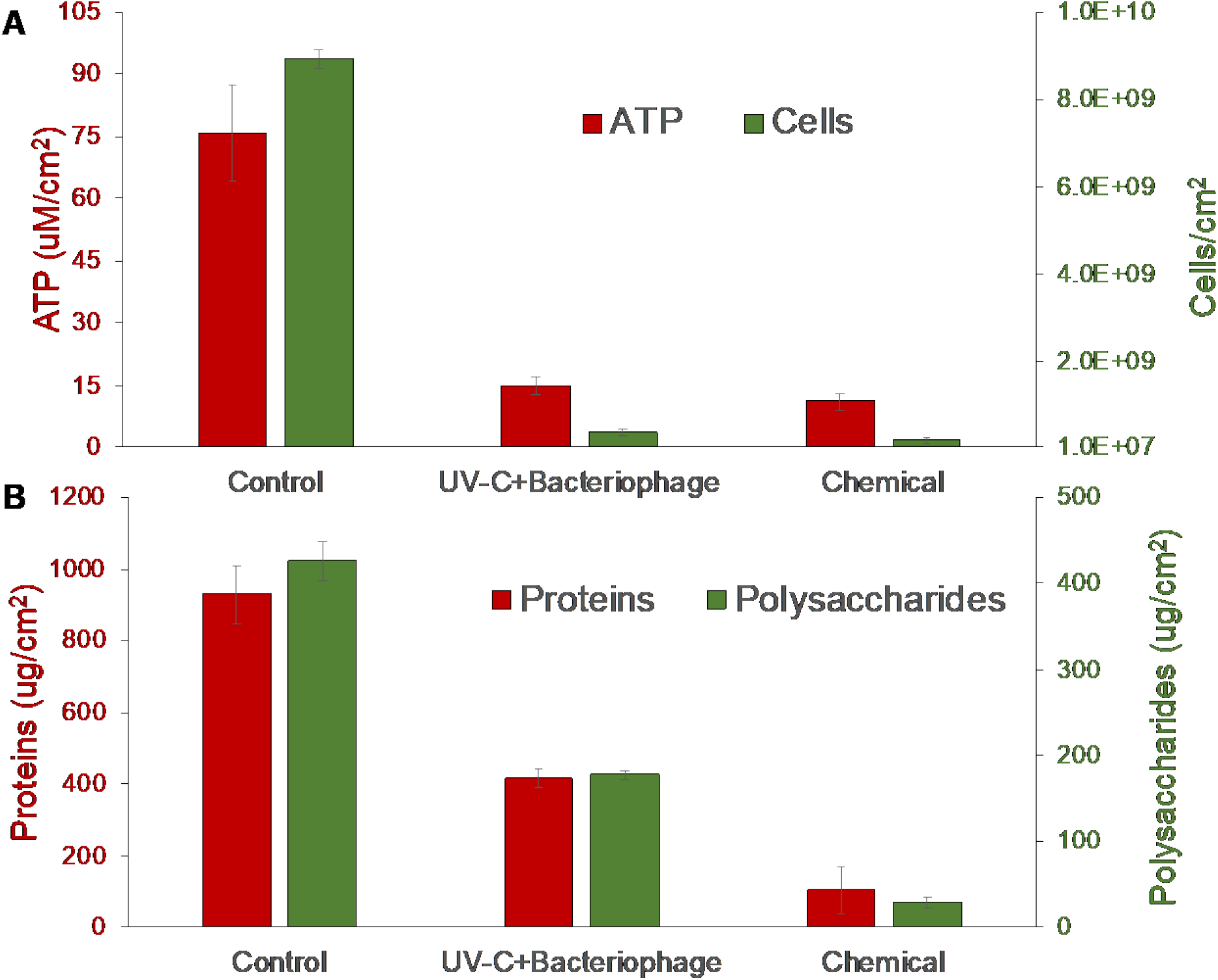
ATP concentration and cells number (A), protein and polysaccharides concentration (B) in membrane biofilm in control, UV-C + bacteriophages cocktail treatment and chemicals addition. The biofilm was collected at a TMP of 40 kPa. Error bars indicate standard deviation among the three replicates.

## DISCUSSION

In earlier studies, bacteriophage infection was used successfully against single-species biofilms on membranes. To exemplify, the addition of bacteriophages to membrane-associated single species biofilm led to a 40% reduction of attached cells (22) and 70% flux recovery (23). Bacteriophages were also embedded on an ultrafiltration membrane to reduce *Escherichia coli* biofilm (24). However, such demonstrations do not represent the effectiveness of bacteriophages against membrane biofouling arising from a diverse variety of bacteria. We observed that bacteriophages infection alone cannot consistently reduce membrane biofouling caused by the diverse microbial community. For instance, when membrane biofilm was harvested at a TMP of 40 and 60 kPa, bacteriophage cocktail application resulted in only slight reduction of bacterial cells, ATP concentration (Figure 1) and alive and dead cell ratio (Table 1). Furthermore, when membranes harvested at 20 kPa (i.e., at early stages of fouling) were treated with bacteriophage cocktail, the bacteriophages were not able to propagate (Figure S2) to numbers that would significantly impact membrane biofouling (Figure 1). This poor effect may be due to a low number of viable target bacteria present within the membrane biofilm. It was previously observed that 10^6^ CFU/mL of the target bacteria is necessary to initiate phage infection (33). The targets of the three bacteriophages from the *Podoviridae* family isolated in this study (Figure S1) were three species of the *Acinetobacter* genus, namely *A. modestus*, *A. junii* and *A. seohaensis*. Although *Acinetobacter spp*. was considered a core component of biofilm structure in AnMBR, and with a relative abundance ranging from 3 to 4 % (9), it is likely that the total number of *Acinetobacter spp.* bacteria within a young biofilm was not enough to initiate an effective infection.

To improve the effect of bacteriophages on membrane-associated biofilm, we proposed the combination of bacteriophages and UV-C irradiation as a membrane cleaning approach. UV-C interacts with bacteria in the biofilm and causes DNA damage due to pyrimidine dimerization. Bacteria exposed to UV-C lose their ability to replicate, which leads to cell death (34). In earlier studies, UV-C irradiation was used as pretreatment to reduce biofouling in wastewater treatment both alone or in combination with chlorine (27, 35). Moreover, when it was applied as anti-biofilm agent in previous studies (36, 37), the cell log reduction was in agreement with that observed in our study. However, UV-C alone is relatively ineffective against gram-positive and spore formers that may be present in the membrane biofilm. Outer membrane of gram-positive bacteria provides a certain level of resistance by blocking UV penetration inside the cell (38). In addition, spores formed from gram-positive bacteria are 5 to 50 times more resistant to UV-C irradiation compared with growing cells (39). Such resistance to UV-C may explain for the increase in relative abundances of five gram-positive bacteria with UV-C treatment (Table 2).

This limitation of UV-C is, however, addressed by pairing it with bacteriophages. Although the bacteriophages isolated in this study were specific to *Acinetobacter* spp., other genera were also detrimentally affected in their relative abundance (Table 2). This may be due to a variety of phage proteins such as holins, endolysin and spanins that can act on other microorganisms, particularly Gram-positive bacteria (40). Some bacteriophages also encode for enzymes that damage the EPS, which would disrupt the biofilm matrix (20). Pairing UV-C with bacteriophages can also trigger the latter into lytic mode (41, 42), as is exemplified from the reduced membrane cleaning efficiency which bacteriophages introduced independently from UV-C (Figure S3).

We observed that both the bacteriophages and UV-C acted together synergistically to disrupt the biofilm matrix and to enhance cell removal compared with either of the individual treatment. The mechanism behind this synergism is speculated to be achieved via two steps. First, UV-C may act on the whole bacterial community by killing the bacteria and loosening the biofilm matrix. This biofilm dislodging effect was observed from the OCT analysis (43, 44), in which cavities in the biofilm matrix were characterized by a lower biofilm coverage and a reduction of thickness and biovolume compared with the control (Figure 3). Second, by loosening the biofilm matrix, bacteriophages are able to penetrate deeper into the biofilm matrix to infect those embedded nearer to the membrane surface. Our results showed that this additive effect was particularly optimal when membranes were subcritically fouled at 40 kPa (Figure 1). At this TMP value, it is likely that the presence of a minimum target bacterial density and a biofilm matrix that was not extremely thick and compact would provide the optimal conditions for UV-C action and bacteriophages infection. In addition, membrane biofilm harvested at 40 kPa showed the highest cell activity compared with the other biofouling rates (Figure 1B). High host activity is a crucial trait that favors phage infection since bacteriophages use host machinery to replicate and to propagate inside the cell (42).

Despite UV-C + bacteriophage effectively reducing bacterial cell numbers and activity within the biofilm (Figure 1 and Table 1), the approach was less effective in reducing protein and polysaccharides concentration in biofilm matrix when compared with the sodium hypochlorite and citric acid cleaning (Figure 4B). Sodium hypochlorite and citric acid are commonly used to clean fouled membranes. Sodium hypochlorite is a strong oxidant that modifies the cell membrane permeability by reacting with phospholipids (45). Citric acid chelates inorganic minerals and disrupts the stability of the biofilm matrix (46, 47). Therefore, while chemical cleaning is directed to the removal of biofilm EPS and irreversible foulant layer, UV-C and bacteriophages may be more useful against reversible foulants formed during the subcritical stages of fouling (e.g. at TMP of 40 kPa).

Considering that bacteriophages are already capable of reducing biofilm thickness and biovolume after 1 h of application (Table S2), we envision that UV-C + bacteriophage as a membrane cleaning method can possibly be carried out in a semi-continuous mode with minimal downtime to the operation process. This strategy could potentially delay the occurrence of membrane fouling by removing the reversible fouling layers developed on membranes, in turn reducing the frequencies and amount of chemicals needed throughout the course of operation.

## MATERIALS AND METHODS

### Bacteriophages isolation and characterization

Bacteriophages targeting three species from the *Acinetobacter* genus, namely *A. modestus, A junii* and *A. seohaensis* were isolated from the influent of the KAUST wastewater treatment plant as described before (21) and propagated by the double layer method (48). A detailed description of bacteriophages isolation can be found in supporting information, section “Bacteriophages isolation”.

Phage morphology was characterized by transmission electron microscopy (TEM) (Tecnai Spirit TWIN, FEI) operated at 120 kV and equipped with an ORIUS SC1000 camera (Gaitan). To obtain TEM photos, the three bacteriophages in suspension were first fixed with 2.5% v/v glutaraldehyde, and then placed on carbon-coated copper grids before being negatively stained with 1% w/v uranyl acetate.

### Reactor set up and operations

AnMBR configuration (Figure S5A) used was similar to the one in earlier study (49). A 2 L reactor, operated without air sparging, was filled with ceramic rings to support biofilm establishment and it was inoculated with the same seed sludge described elsewhere (7). Synthetic wastewater with a chemical oxygen demand (COD) of 750 mg/L was fed into the reactor with a hydraulic retention time of 18.5 h. The reactor was operated at pH 7 and at 35 °C and no sludge was wasted during the entire study. The reactor was connected to three PVDF microfiltration membrane (0.3 μm nominal pore size, GE Osmonics, Minnetonka, MN) modules in parallel. The three membranes were operated in cross-flow mode with stable flux that ranged from 6 to 7 L/m^2^/h (LMH). Biogas was used to scour the membrane surface. Transmembrane pressure (TMP) was recorded every day by a pressure gauge connected to each membrane module. Following the indication of a previous study (49), membranes were harvested and analyzed at three different TMP values, i.e. 20, 40 and 60 kPa, which represent increasing biofouling extent (Figure S5B). Three biological replicates of fouled membranes were harvested at each TMP values.

### Membrane harvesting and treatments application

Harvested membranes were processed for analysis. The first 3 cm at each end of the membranes were discarded because biofilm formation was not homogenous at those locations. The rest of the membrane, with homogenous biofilm, was cut into four pieces with dimensions 2 by 2 cm. These pieces were individually placed in small sterile petri dish that contained 1X phosphate buffer solution (PBS), and subsequently incubated at room temperature for 6 h. Before the incubation, three types of treatments were applied, namely (i) bacteriophages cocktail treatment, named “bacteriophage”, with a total multiplicity of infection (MOI) of 1; (ii) UV-C treatment, by exposing the biofilm to 100 mJ/cm^2^ twice, at 3 and 6 h from the start of the incubation; (iii) a combination of the three *Acinetobacter* bacteriophages at an MOI of 1 and UV-C irradiation exposing the biofilm to 100 mJ/cm^2^ twice, at 3 and 6 h from the start of the incubation (this treatment was named “UV-C + bacteriophage”). A last treatment involves UV-C being applied before bacteriophages addition in an independent manner (this treatment was named “independent UV-C + bacteriophage” and more details are provided in supporting information, section “Effect of UV-C and bacteriophages application order”). Lastly, one piece of membrane was soaked in 1X PBS for 6 h to represent control. The bacteriophages MOI was set based on the relative abundances of *Acinetobacter* spp. observed in a previous study (9).

At the end of the 6 h, 1 mL of the solution in which the membrane was soaked was aliquot for PFU counting as described in the next section. Afterwards, the membrane piece was removed from the solution, washed and placed in fresh 1X PBS. Attached biofilm was sonicated for 5 min using a Q500 sonicator (Qsonica, Newton, CT, US) at 25% amplitude with 5 s pulsating step. The rest of the biofilm was scraped using a sterile inoculation loop.

### Bacteriophages plaque counts after the treatments

The 1 mL aliquot was filtered through 0.22 μm syringe filter to remove remaining biomass. Several dilutions, ranging from 10^−1^ to 10^−4^ fold were performed in sodium magnesium buffer (5.8 g/L NaCl, 0.975 g/L MgSO_4_, 50 mL/L 50 mM Tris-Cl at pH 7.5), and 10 μL of the diluted suspension was mixed individually with 100 μL of *A. junii*, *A. modestus* and *A. seohaensis* cultures. The presence of plaques was identified with the double layer method (48) and it was compared with the initial amount of PFU/mL of each bacteriophage spiked for the treatments (5.4 × 10^7^ PFU/mL of *A. junii* bacteriophage, 6.9 × 10^7^ PFU/mL of *A. modestus* bacteriophage and 2.1 × 10^6^ PFU/mL of *A. seohaensis* bacteriophage).

### Biofilm characterization

Biofilm structure was characterized by enumerating the total cells, the proportion of cells with intact cell wall membranes, adenosine triphosphate (ATP) concentration and EPS concentration.

To enumerate cells, membrane biofilm suspension was diluted by 10^4^, stained with SYBR^®^ green (Thermo Fisher Scientific, Walthman, MA, US) for 15 min at 37 °C and counted using a BD Accurri flow cytometer (BD Bioscience, Franklin Lakes, NJ). The ratio between alive and dead cells was calculated by staining the same diluted biofilm suspension using the LIVE/DEAD^®^ BacLight™ Bacterial Viability and Counting kit (Thermo Fisher Scientific, Walthman, MA, US) for 15 min in the dark at room temperature. Alive and dead cells were enumerated using a BD Accurri flow cytometer (BD Bioscience, Franklin Lakes, NJ). ATP concentration was measured using a Celsis ATP reagent kit and an Advance luminometer (Celsis, Westminster, United Kingdom).

For EPS analysis, both polysaccharides and proteins concentration were measured as described in a previous study (50). Firstly, 2 mL of biofilm suspension was filtered through 0.22 μm syringe filter to quantify only the filtrate (i.e., dissolved constituents). For each piece of cut membrane, total protein concentration was measured in triplicates by Total Protein kit (Sigma-Aldrich, St. Luis, MO, US) using bovine serum albumin (BSA) as standard (Sigma-Aldrich, St. Luis, MO, US). Finally, polysaccharides were quantified by the phenol-sulfuric method (51), using glucose as standard (Sigma-Aldrich, St. Luis, MO, US). Briefly, 1 mL of each sample (including different concentrations of glucose standard) was mixed with 1 mL of 5% v/v phenol solution and with 5 mL 98% sulfuric acid (Sigma-Aldrich, St. Luis, MO, US). The mixture was incubated at room temperature for 20 min before measuring optical density at 490 nm.

### RNA extraction and biofilm microbial community analysis

To analyze the composition of the active biofilm bacterial community, an aliquot of 3 mL of suspended biofilm was used to extract the RNA that was then reverse transcribed into first-strand complementary DNA. 16S rRNA gene amplicon sequencing was performed to assess the influence of the different treatment on the membrane biofilm community as described before (52). More details can be found in supporting information, section “RNA extraction and amplicon sequencing analysis”. All high-throughput sequencing files were deposited in the archive of the European Nucleotide Archive under study accession number PRJEB38595.

### Optical coherence tomography analysis

A spectral domain optical coherence tomography SD-OCT system device (Ganymede I, Thorlabs GmbH, Dachau, Germany) provided with LSM03 scan lens was employed to non-invasively evaluate the effect of the treatments on the biomass developed on the membrane surface. The OCT employs backscattered light to acquire cross-sectional scans of membrane. A membrane module connected to the anaerobic reactor was operated to reach a TMP of 40 kPa. This TMP value was chosen because it resulted in the best biofilm removal for the three treatments compared with the control. The membrane was therefore harvested at TMP of 40 kPa and cut to four pieces for individual application of the three treatments (i.e., bacteriophage, UV-C, UV-C+ bacteriophage), with the remaining portion used as non-treated control. At the end of the 6 h treatment, the membrane pieces in the petri dish were positioned under the OCT probe to assess the efficiency of each treatment in removing the biomass from the membrane surface. 3D images for each analysis were obtained to show and visualize the biofouling reduction. In addition, a time-series analysis was performed to monitor the effect of the bacteriophage treatment alone by fixing the membrane coupons under the OCT probe.

The biomass morphology descriptors, namely biovolume, biofilm coverage, average thickness and the relative roughness were calculated using the equation reported in literature (43, 44), by analyzing 500 scan images for the 3D scan. More details can be found in supporting information, section “OCT images analysis”.

### Comparison with chemical membrane cleaning

We further compared the UV-C + bacteriophage treatment to the conventional cleaning process. Briefly, a membrane module connected to the anaerobic reactor was harvested at TMP of 40 kPa, and cut into three pieces of dimension 2 by 2 cm. The pieces were individually were placed in a small petri dish filled with 1X PBS. One of these pieces was exposed to the same mixed treatment described above (UV-C + bacteriophages); the second piece was soaked in a mixture of 0.1 M citric acid and 6% sodium hypochlorite solution; the last piece of membrane was used as control submerged in 1X PBS.

After 6 h, the membrane was removed from the solution and the biofilm was resuspended in fresh 1X PBS as described before. The experiment was performed in triplicates. The effect of the two different treatments compared to the control was examined with the same analysis described in section “Biofilm characterization”.

### Statistical analysis

Statistical differences for the parameters at different conditions were evaluated through one-way ANOVA with significance level set at 95% confidence level (p < 0.05).

## ACKNOWLEDGEMENTS

This study was supported by KAUST Center Applied Research Funding FCC/1/1971-32-01 and Center of Excellence for NEOM Research flagship projects REI/1/4178-03-01 awarded to P.Y.-H.

## SUPPORTING INFORMATION

### Bacteriophages isolation

Briefly, 50 mL of freshly collected influent was centrifuged for 20 min at 8,500 *g.* The supernatant was filtered through 0.22 μm cellulose acetate syringe filter (VWR, Radnor, PA) and then mixed with 50 mL of LB broth and 50 mL of exponentially growing cultures of each *Acinetobacter* isolates mentioned above. This mixture was incubated at 37 °C for 24 h before cells lysis by injecting 1% v/v chloroform. After 2 h incubation at room temperature and 100 rpm agitation, the mixture was centrifuged at 8,500 *g* for 30 min and the supernatant was filtered to remove remaining bacterial cells. Aliquots of 100 μL of the filtered supernatant were used to identify the presence of plaques using the double layer method. Single plaques were then isolated with inoculating loops, transferred to SM buffer and filtered to remove bacterial cells. After propagating several times, the plaque forming units (PFU/mL) were determined for each of the three bacteriophages.

### RNA extraction and amplicon sequencing analysis

RNA extraction was performed with the RNeasy^®^ midi kit (Qiagen, Hilden, Germany), including a DNase treatment step. RNA concentration was quantified with Invitrogen RNA HS Qubit^®^ 2.0 assay kit (Thermo Fisher Scientific, Carlsbad, CA, US). RNA was then reverse transcribed into first-strand complementary DNA using the Invitrogen SuperScript™ First-strand Synthesis System (Thermo Fisher Scientific, Carlsbad, CA, US). Starting from this DNA 16S rRNA genes was amplified using the 515F (5’-Illumina overhang-GTG YCA GCM GCC GCG GTA A-3’) and 907R (5’-Illumina overhang-CCC CGY CAA TTC MTT TRA GT-3’) primers pair. DNA amplicons were then submitted to KAUST genomic core lab for unidirectional sequencing on Illumina MiSeq platform. Raw amplicons sequences were sorted on a Phred score > 30, before primers, adapters and index sequences removal. UCHIME was used to remove sequence chimeras (53). For each sample, a subset of 100,000 sequences were analyzed and 16S rRNA gene sequences were annotated for their taxonomical assignments using RDP classifier at a 95% confidence level (54). 16S rRNA gene copy number adjustment for each taxonomical unit was also performed. To assess the impact of the different treatment on *Acinetobacter*-related taxa and, in general on the whole biofilm structure, the bacterial community was characterized in terms of relative abundance of genus level.

### OCT images analysis

3-D cross-sectional scans of 800 × 800 × 516 pixels (width × length × depth), corresponding to a volume of 8.0 × 8.0 × 1.4 mm were acquired. The image analysis on the OCT scans was performed through a customized MATLAB code. The data sets corresponding to the 3D OCT cross sectional scans were visualized by using Avizo software. The membrane and biofilm were defined using Avizo’s segmentation editor. The 3D rendering volume was generated to visualize over time the effect of the bacteriophage treatment on the biofilm deposited on the membrane surface.

### Effect of UV-C and bacteriophages application order

For all the experiments in this study, the combination of UV-C and bacteriophages cocktail was applied by infecting the biofilm with phages at the beginning of the treatment, while UV-C was irradiated at 100 mJ/cm^2^ after 3 and 6 h (UV-C + Bacteriophages). To assess if the order of applying UV-C and bacteriophages would affect the treatment efficacy, a separate experiment was made in which UV-C was applied before bacteriophages was spiked. In this treatment, bacteriophages were not exposed to UV-C. This treatment was named “Independent UV-C + bacteriophages”. To elucidate, membrane biofilm was harvested at a TMP of 60 kPa and processed as described in section “Membrane harvesting and treatments application”. One fouled membrane piece of dimensions 2 by 2 cm was treated with the usual UV-C + bacteriophages treatment described in the section “Membrane harvesting and treatments application”, while another piece with the same dimension was first irradiated with the same UV-C intensity (200 mJ/cm^2^) and subsequently infected with the bacteriophages cocktail for 6 h in the absence of UV-C. A non-treated fouled membrane piece was used as control. Antifouling effect was characterized in terms of total cells number and proteins and polysaccharides concentration. The same analyses were conducted as described in section “Biofilm characterization” to assess the efficacy of these treatment.

---

**Figure S1.**
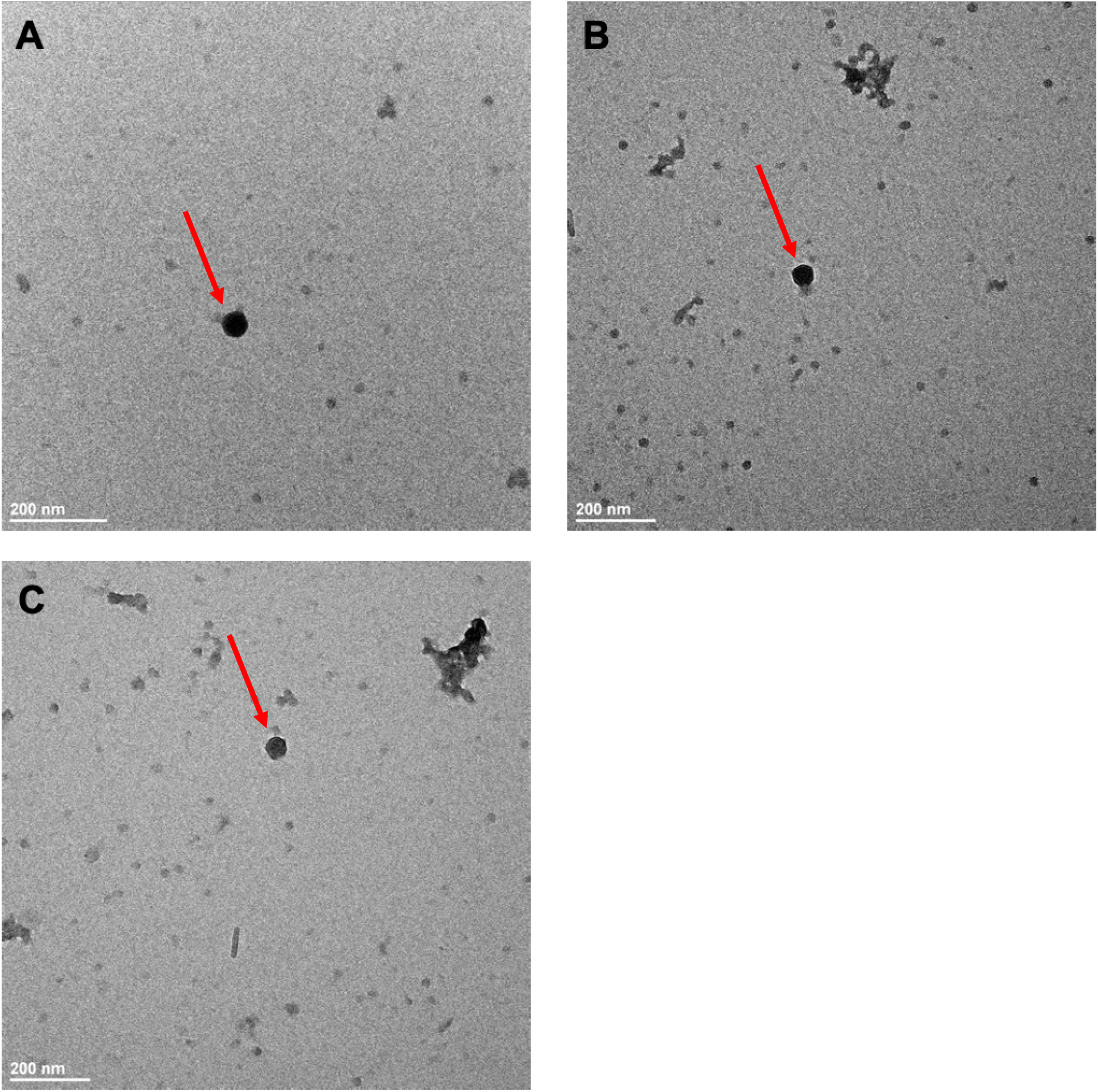
Transmission electron microscopy images of the isolated A*cinetobacter junii* (A), *Acinetobacter modestus* (B) *and Acinetobacter seohaensis* (C) bacteriophages

**Figure S2.**
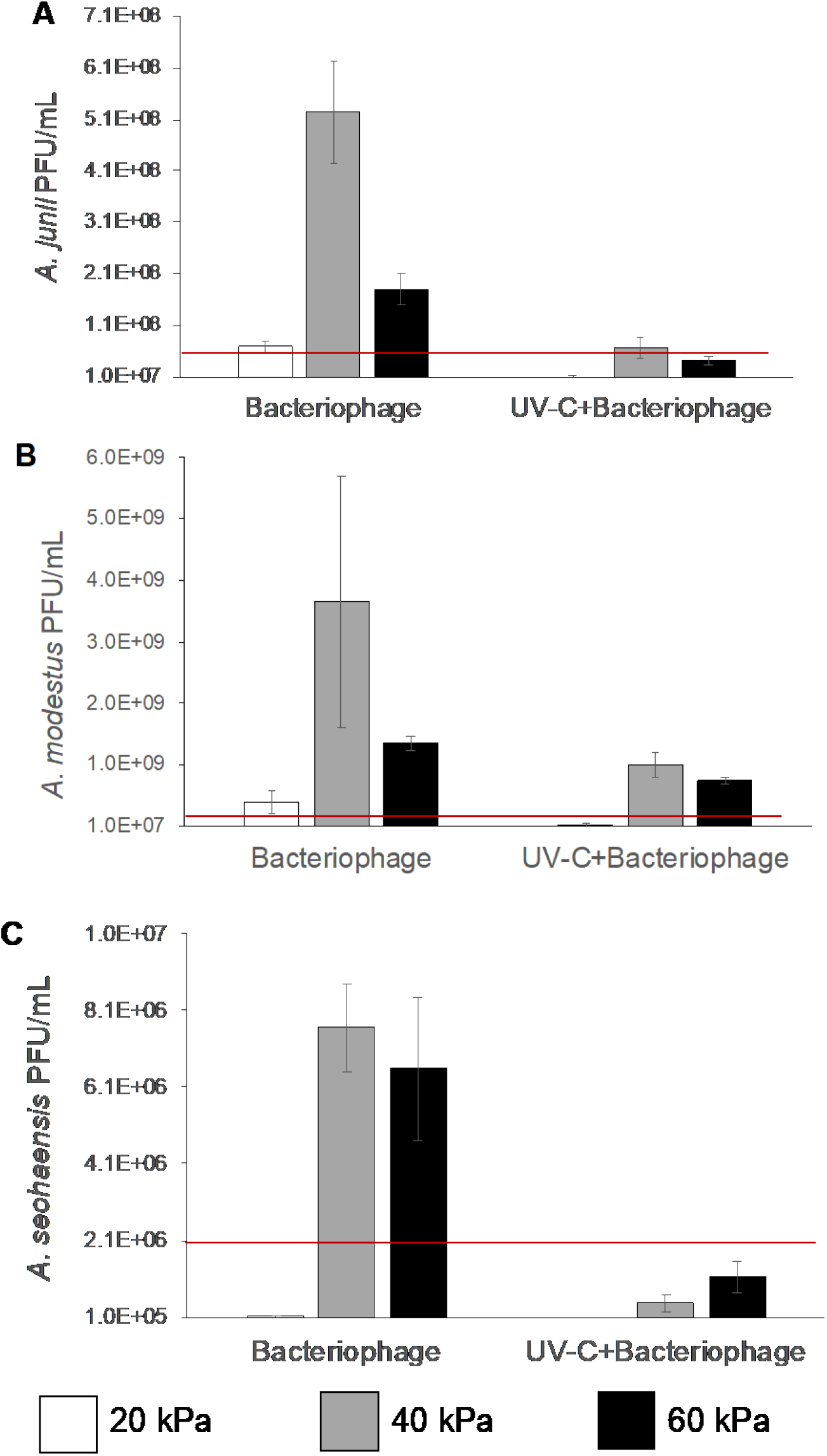
PFU/mL of *A. junii* (A), *A. modestus* (B) and *A. seohaensis* (C) bacteriophages recovered from the soaking solution after membrane biofilm treatment at different biofouling degree. The red line represents the PFU/mL of bacteriophages spiked at the beginning of the treatment. Error bars indicate standard deviation among the three replicates.

**Figure S3.**
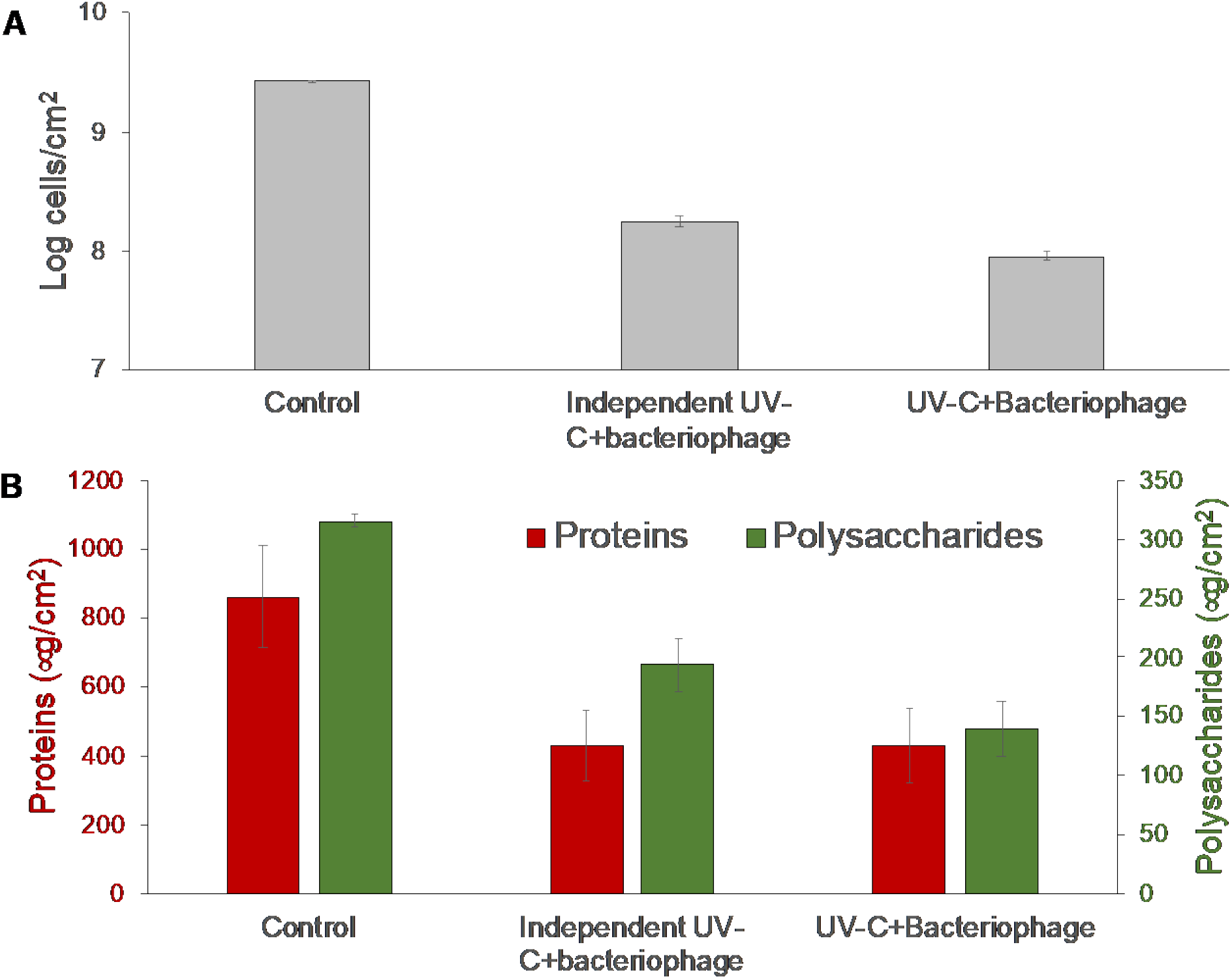
Log of the total cells number (A) and proteins and polysaccharides concentration (B) for the control, Independent UV-C + bacteriophage treatment, in which UV-C was irradiated before bacteriophages application and UV-C + bacteriophage treatment, in which the two agents were utilized in combination.

**Figure S4.**
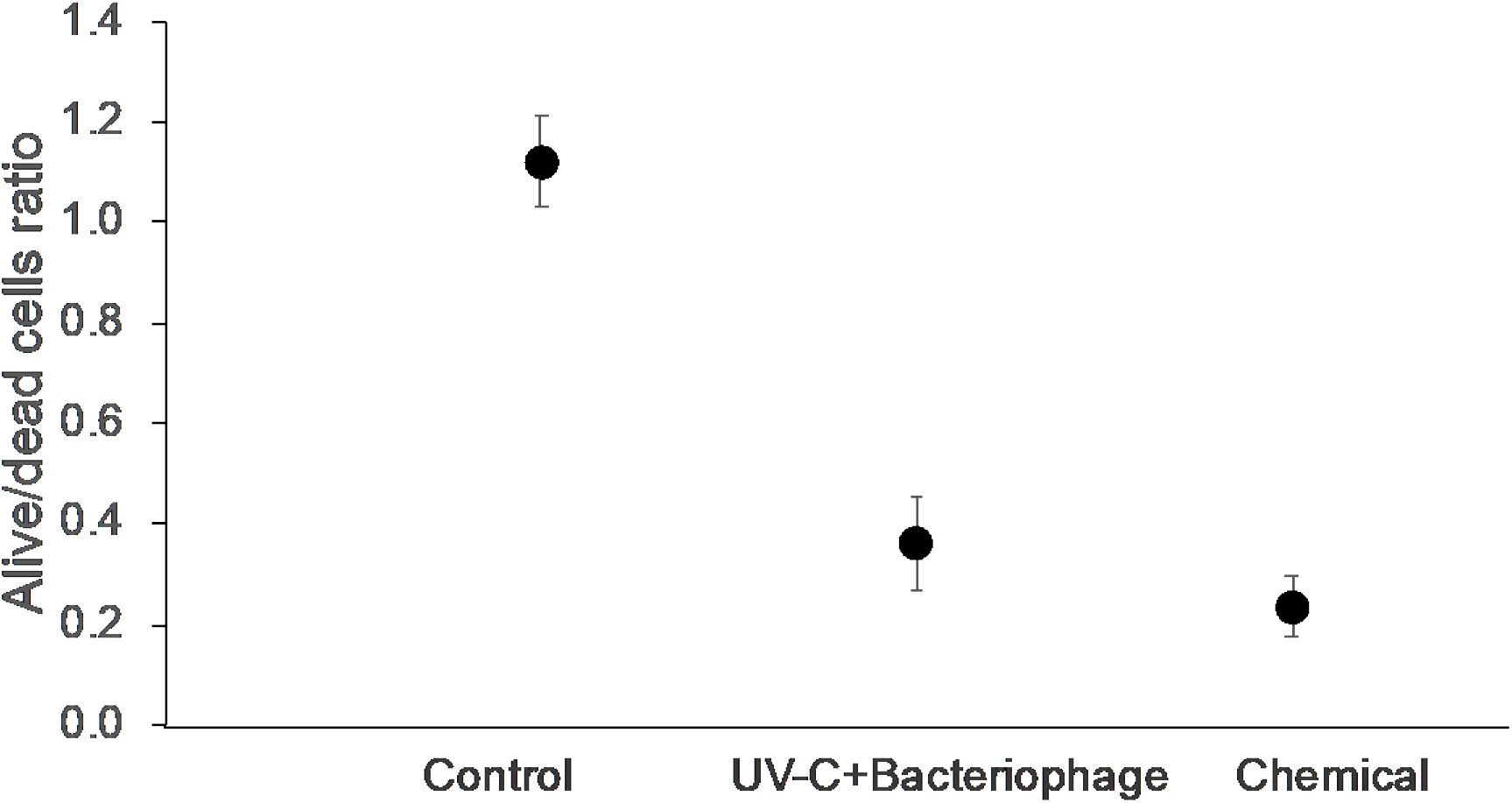
Ratio between alive and dead cells in membrane biofilm in control, UV-C + bacteriophage cocktail treatment and chemicals treatment. The biofilm was collected at a TMP of 40 kPa. Error bars indicate standard deviation among the three replicates.

**Figure S5.**
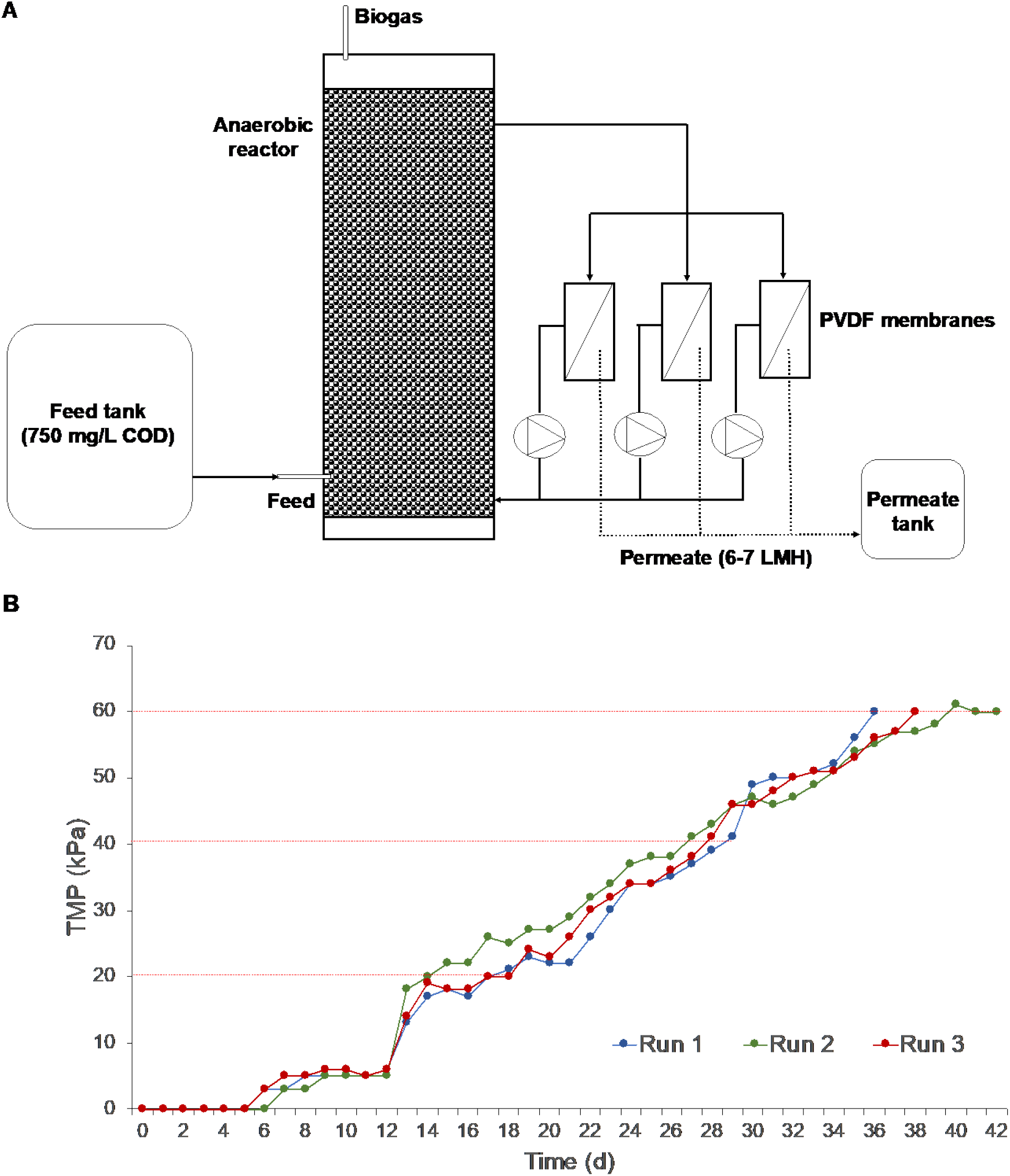
Scheme of the anaerobic reactor and the three parallel cross-flow membranes used in this study (A). Transmembrane pressure profiles of PVDF membranes. The three biological replicates are indicated as run 1, run 2 and run 3. Red dashed lines indicate the biofouling rates (20, 40 and 60 kPa) at which the membranes were harvested (B)

**Table S1.** Characterization of the membrane biofilm after different treatments and at different biofouling rates. The data were obtained from OCT analysis and depict the values of 500 scan images for a total area of 8 × 8 × 1.4 mm.

| Condition | Thickness ($\mu\text{m}$) | Roughness | Biovolume ($\text{mm}^3/\text{cm}^2$) | Membrane coverage % |
| --- | --- | --- | --- | --- |
| Control | $43 \pm 1$ | $0.11 \pm 0.02$ | $4.3 \pm 0.1$ | 99.5 |
| Bacteriophage | $39 \pm 4$ | $0.21 \pm 0.07$ | $3.4 \pm 0.4$ | 86.7 |
| UV | $38 \pm 5$ | $0.18 \pm 0.12$ | $3.4 \pm 0.5$ | 88.5 |
| UV-C + Bacteriophage | $38 \pm 3$ | $0.17 \pm 0.05$ | $3.6 \pm 0.3$ | 89.5 |

**Table S2.** Changes in membrane biofilm with time after *Acinetobacter* bacteriophages cocktail application. The data were obtained from the values of 500 scan images for a total volume of 8.0 × 8.0 × 1.4 mm.

| Time (h) | Thickness ($\mu\text{m}$) | Roughness | Biovolume ($\text{mm}^3/\text{cm}^2$) |
| --- | --- | --- | --- |
| 0 | $42 \pm 4$ | $0.13 \pm 0.02$ | $4.2 \pm 0.2$ |
| 1 | $34 \pm 2$ | $0.12 \pm 0.01$ | $3.4 \pm 0.2$ |
| 3 | $34 \pm 2$ | $0.12 \pm 0.02$ | $3.4 \pm 0.2$ |
| 4.5 | $30 \pm 2$ | $0.22 \pm 0.11$ | $3.0 \pm 0.2$ |
| 6 | $30 \pm 2$ | $0.24 \pm 0.12$ | $3.0 \pm 0.2$ |

## REFERENCES

1. A. L. Smith, L. B. Stadler, N. G. Love, S. J. Skerlos, L. Raskin, Perspectives on anaerobic membrane bioreactor treatment of domestic wastewater: a critical review. Bioresource technology 122, 149–159 (2012).

2. A. L. Prieto, H. Futselaar, P. N. Lens, R. Bair, D. H. Yeh, Development and start up of a gas-lift anaerobic membrane bioreactor (Gl-AnMBR) for conversion of sewage to energy, water and nutrients. Journal of membrane science 441, 158–167 (2013).

3. D. Rosso, L. E. Larson, M. K. Stenstrom, Aeration of large-scale municipal wastewater treatment plants: state of the art. Water Science and Technology 57, 973–978 (2008).

4. B.-Q. Liao, J. T. Kraemer, D. M. Bagley, Anaerobic membrane bioreactors: applications and research directions. Critical Reviews in Environmental Science and Technology 36, 489–530 (2006).

5. M. Maaz et al., Anaerobic membrane bioreactors for wastewater treatment: Novel configurations, fouling control and energy considerations. Bioresource technology (2019).

6. M. Harb, P.-Y. Hong, Anaerobic membrane bioreactor effluent reuse: A review of microbial safety concerns. Fermentation 3, 39 (2017).

7. Y. Xiong, M. Harb, P.-Y. Hong, Characterization of biofoulants illustrates different membrane fouling mechanisms for aerobic and anaerobic membrane bioreactors. Separation and Purification Technology 157, 192–202 (2016).

8. P. Xu, J. E. Drewes, T.-U. Kim, C. Bellona, G. Amy, Effect of membrane fouling on transport of organic contaminants in NF/RO membrane applications. Journal of Membrane Science 279, 165–175 (2006).

9. H. Cheng, D. Cheng, J. Mao, T. Lu, P.-Y. Hong, Identification and characterization of core sludge and biofilm microbiota in anaerobic membrane bioreactors. Environment international 133, 105165 (2019).

10. D. P. Thanu, M. Zhao, Z. Han, M. Keswani, “Fundamentals and Applications of Sonic Technology” in Developments in Surface Contamination and Cleaning: Applications of Cleaning Techniques. (Elsevier, 2019), pp. 1–48.

11. H. Lin, J. Chen, F. Wang, L. Ding, H. Hong, Feasibility evaluation of submerged anaerobic membrane bioreactor for municipal secondary wastewater treatment. Desalination 280, 120–126 (2011).

12. H. Sanawar, Y. Xiong, A. Alam, J.-P. Croue, P.-Y. Hong, Chlorination or monochloramination: Balancing the regulated trihalomethane formation and microbial inactivation in marine aquaculture waters. Aquaculture 480, 94–102 (2017).

13. J. Le Roux, N. Nada, M. T. Khan, J.-P. Croué, Tracing disinfection byproducts in full-scale desalination plants. Desalination 359, 141–148 (2015).

14. M. R. Jumat, M. F. Haroon, N. Al-Jassim, H. Cheng, P.-Y. Hong, An increase of abundance and transcriptional activity for Acinetobacter junii post wastewater treatment. Water 10, 436 (2018).

15. Y. Xiong, Y. Liu, Biological control of microbial attachment: a promising alternative for mitigating membrane biofouling. Applied microbiology and biotechnology 86, 825–837 (2010).

16. L. Malaeb, P. Le-Clech, J. S. Vrouwenvelder, G. M. Ayoub, P. E. Saikaly, Do biological-based strategies hold promise to biofouling control in MBRs? Water research 47, 5447–5463 (2013).

17. K. S. Liao, S. M. Lehman, D. J. Tweardy, R. M. Donlan, B. W. Trautner, Bacteriophages are synergistic with bacterial interference for the prevention of Pseudomonas aeruginosa biofilm formation on urinary catheters. Journal of applied microbiology 113, 1530–1539 (2012).

18. D. Alemayehu et al., Bacteriophages φMR299–2 and φNH-4 can eliminate Pseudomonas aeruginosa in the murine lung and on cystic fibrosis lung airway cells. MBio 3, e00029–00012 (2012).

19. D. P. Pires, D. V. Boas, S. Sillankorva, J. Azeredo, Phage therapy: a step forward in the treatment of Pseudomonas aeruginosa infections. Journal of virology 89, 7449–7456 (2015).

20. D. R. Harper et al., Bacteriophages and biofilms. Antibiotics 3, 270–284 (2014).

21. G. Scarascia, S. A. Yap, A. H. Kaksonen, P.-Y. Hong, Bacteriophage infectivity against Pseudomonas aeruginosa in saline conditions. Frontiers in microbiology 9, 875 (2018).

22. G. Goldman, J. Starosvetsky, R. Armon, Inhibition of biofilm formation on UF membrane by use of specific bacteriophages. Journal of Membrane Science 342, 145–152 (2009).

23. A. S. Bhattacharjee, J. Choi, A. M. Motlagh, S. T. Mukherji, R. Goel, Bacteriophage therapy for membrane biofouling in membrane bioreactors and antibiotic - resistant bacterial biofilms. Biotechnology and bioengineering 112, 1644–1654 (2015).

24. W. Ma, M. Panecka, N. Tufenkji, M. S. Rahaman, Bacteriophage-based strategies for biofouling control in ultrafiltration: in situ biofouling mitigation, biocidal additives and biofilm cleanser. Journal of colloid and interface science 523, 254–265 (2018).

25. Y. Zhang, Z. Hu, Combined treatment of Pseudomonas aeruginosa biofilms with bacteriophages and chlorine. Biotechnology and bioengineering 110, 286–295 (2013).

26. R. Gehr, M. Wagner, P. Veerasubramanian, P. Payment, Disinfection efficiency of peracetic acid, UV and ozone after enhanced primary treatment of municipal wastewater. Water research 37, 4573–4586 (2003).

27. A. Benito, G. Garcia, R. Gonzalez-Olmos, Fouling reduction by UV-based pretreatment in hollow fiber ultrafiltration membranes for urban wastewater reuse. Journal of Membrane Science 536, 141–147 (2017).

28. N. Pozos, K. Scow, S. Wuertz, J. Darby, UV disinfection in a model distribution system:: biofilm growth and microbial community. Water Research 38, 3083–3091 (2004).

29. Y. Zhu et al., Characterization of biofilm and corrosion of cast iron pipes in drinking water distribution system with UV/Cl2 disinfection. Water research 60, 174–181 (2014).

30. N. Trun, J. Trempy, Fundamental bacterial genetics (John Wiley & Sons, 2009).

31. H.-W. Ackermann, Classification of bacteriophages. The bacteriophages 635, 8–16 (2006).

32. L. Fortunato, A. F. Lamprea, T. Leiknes, Evaluation of membrane fouling mitigation strategies in an algal membrane photobioreactor (AMPBR) treating secondary wastewater effluent. Science of The Total Environment 708, 134548 (2020).

33. S. Chibani-Chennoufi, A. Bruttin, M.-L. Dillmann, H. Brüssow, Phage-host interaction: an ecological perspective. Journal of bacteriology 186, 3677–3686 (2004).

34. N. Goosen, G. F. Moolenaar, Repair of UV damage in bacteria. DNA repair 7, 353–379 (2008).

35. J. Xing et al., Application of low-dosage UV/chlorine pre-oxidation for mitigating ultrafiltration (UF) membrane fouling in natural surface water treatment. Chemical Engineering Journal 344, 62–70 (2018).

36. Y. M. Bae, S. Y. Lee, Inhibitory effects of UV treatment and a combination of UV and dry heat against pathogens on stainless steel and polypropylene surfaces. Journal of food science 77, M61–M64 (2012).

37. C. Hadjok, G. Mittal, K. Warriner, Inactivation of human pathogens and spoilage bacteria on the surface and internalized within fresh produce by using a combination of ultraviolet light and hydrogen peroxide. Journal of applied microbiology 104, 1014–1024 (2008).

38. P. D. Williams, S. L. Eichstadt, T. A. Kokjohn, E. L. Martin, Effects of ultraviolet radiation on the gram-positive marine bacterium Microbacterium maritypicum. Current microbiology 55, 1–7 (2007).

39. P. Setlow, Resistance of spores of Bacillus species to ultraviolet light. Environmental and molecular mutagenesis 38, 97–104 (2001).

40. R. Young, Phage lysis: do we have the hole story yet? Current opinion in microbiology 16, 790–797 (2013).

41. N. Al-Jassim, D. Mantilla-Calderon, G. Scarascia, P.-Y. Hong, Bacteriophages to sensitize a pathogenic New Delhi metallo β-lactamase-positive Escherichia coli to solar disinfection. Environmental science & technology 52, 14331–14341 (2018).

42. A. Campbell, The future of bacteriophage biology. Nature Reviews Genetics 4, 471–477 (2003).

43. L. Fortunato et al., Cake layer characterization in activated sludge membrane bioreactors: real-time analysis. Journal of membrane science 578, 163–171 (2019).

44. L. Fortunato, N. Pathak, Z. U. Rehman, H. Shon, T. Leiknes, Real-time monitoring of membrane fouling development during early stages of activated sludge membrane bioreactor operation. Process Safety and Environmental Protection 120, 313–320 (2018).

45. G. Zhen et al., Anaerobic membrane bioreactor towards biowaste biorefinery and chemical energy harvest: Recent progress, membrane fouling and future perspectives. Renewable and Sustainable Energy Reviews 115, 109392 (2019).

46. H. Lee et al., Cleaning strategies for flux recovery of an ultrafiltration membrane fouled by natural organic matter. Water research 35, 3301–3308 (2001).

47. N. Porcelli, S. Judd, Chemical cleaning of potable water membranes: a review. Separation and Purification Technology 71, 137–143 (2010).

48. M. H. Adams, Bacteriophages. Bacteriophages. (1959).

49. H. Cheng, P.-Y. Hong, Removal of antibiotic-resistant bacteria and antibiotic resistance genes affected by varying degrees of fouling on anaerobic microfiltration membranes. Environmental science & technology 51, 12200–12209 (2017).

50. H. Cheng et al., Antibiofilm effect enhanced by modification of 1, 2, 3-triazole and palladium nanoparticles on polysulfone membranes. Scientific reports 6, 24289 (2016).

51. M. Dubois, K. A. Gilles, J. K. Hamilton, P. t. Rebers, F. Smith, Colorimetric method for determination of sugars and related substances. Analytical chemistry 28, 350–356 (1956).

52. G. Scarascia, H. Cheng, M. Harb, P.-Y. Hong, Application of hierarchical oligonucleotide primer extension (HOPE) to assess relative abundances of ammonia-and nitrite-oxidizing bacteria. BMC microbiology 17, 85 (2017).

53. R. C. Edgar, B. J. Haas, J. C. Clemente, C. Quince, R. Knight, UCHIME improves sensitivity and speed of chimera detection. Bioinformatics 27, 2194–2200 (2011).

54. J. R. Cole et al., The Ribosomal Database Project: improved alignments and new tools for rRNA analysis. Nucleic acids research 37, D141–D145 (2009).

